# Establishment of Measles Virus Receptor-expressing Vero Cells Lacking Functional Poliovirus Receptors

**DOI:** 10.1101/2022.07.21.500891

**Authors:** Kenji Someya, Yuko Okemoto-Nakamura, Takako Kurata, Daiki Kanbayashi, Noriko Saito, Masae Itamochi, Noriyuki Otsuki, Kentaro Hanada, Makoto Takeda

## Abstract

Global efforts are underway to eliminate measles and rubella and active viral surveillance is key to achieving this goal. In addition to this, the World Health Organization announced guidelines for handling materials potentially infectious for poliovirus (PV) to minimize the risk of PV reintroduction and achieve PV eradication. To support the global efforts, we have established new PV-non-susceptible cell lines useful for the isolation of measles virus (MeV) and rubella virus (RuV) (Vero ΔPVR1/2 hSLAM+). In the cell lines, MeV and RuV replicated efficiently with no concern about PV replication.

---

To minimize the risk of poliovirus (PV) reintroduction after type-specific eradication, to contain wild PV and to cease oral polio vaccination, the World Health Organization (WHO) announced guidelines for facilities that collect, handle and store materials potentially infectious for PV (1). To propagate measles virus (MeV) or rubella virus (RuV) from clinical specimens containing or potentially containing PV without multiplying PV, we developed Vero ΔPVR1/2 hSLAM+ cells that express the MeV receptor (human signaling lymphocytic activation molecule: hSLAM) (2) but lack functional PV receptors (ΔPVR1/2) (3,4).

Previously, we established a Vero cell line lacking functional PV receptors (Vero ΔPVR1/2 cells [JCRB1840]) (4). Using a retroviral vector system (Cell Biolabs Inc., CA, USA), hSLAM cDNA (NCBI accession number: AY040554) was introduced into the Vero ΔPVR1/2 cells (4) and selected in the presence of blasticidin. We established two blasticidin-resistant cell clones (Vero ΔPVR1/2 hSLAM+ c1 and Vero ΔPVR1/2 hSLAM+ c2). hSLAM expression in these two cell clones was confirmed by flow cytometry (Fig. 1). Vero/hSLAM cells (JCRB1809) (5), established in 2001 and currently used for global surveillance of MeV and RuV (6), were used as a control. To determine whether the two cell clones maintain non-susceptibility to PV, similarly to their parental Vero ΔPVR1/2 cells, the cells were infected with a high titer [multiplicity of infection (MOI) of 10] of Sabin 1 (serotype 1) and Sabin 3 (serotype 3) strains of PV. The cells were successively harvested together with culture supernatants (cell suspensions), and then viral RNA copy numbers were quantified by a real-time PCR assay (7). Viral RNA copy numbers were measured up to 10 days after infection. Both Sabin 1 and Sabin 3 PV strains grew in Vero/hSLAM cells, but there was no increase in viral RNA levels in both Vero ΔPVR1/2 hSLAM+ c1 and Vero ΔPVR1/2 hSLAM+ c2 cells (Fig. 2A, B). We next examined MeV and RuV replication in the two cell clones. The cells were infected with wild-type strain-based recombinant MeV (IC323-EGFP) (8), vaccine strain-based recombinant MeV (AIK-C EGFP) (9) and wild-type strain-based recombinant RuV (rHS717AG1) (10) at a MOI of 0.01. The cell suspensions were harvested successively and viral RNA copy numbers were measured by a real-time PCR assay (11, 12, 13). Compared with Vero/hSLAM cells (5), the two cell clones showed similar levels of MeV RNA production (Fig. 2C, D), whereas, the RuV RNA levels of the two clones were greater than for Vero/hSLAM cells (Fig. 2E).

**Figure 1.**
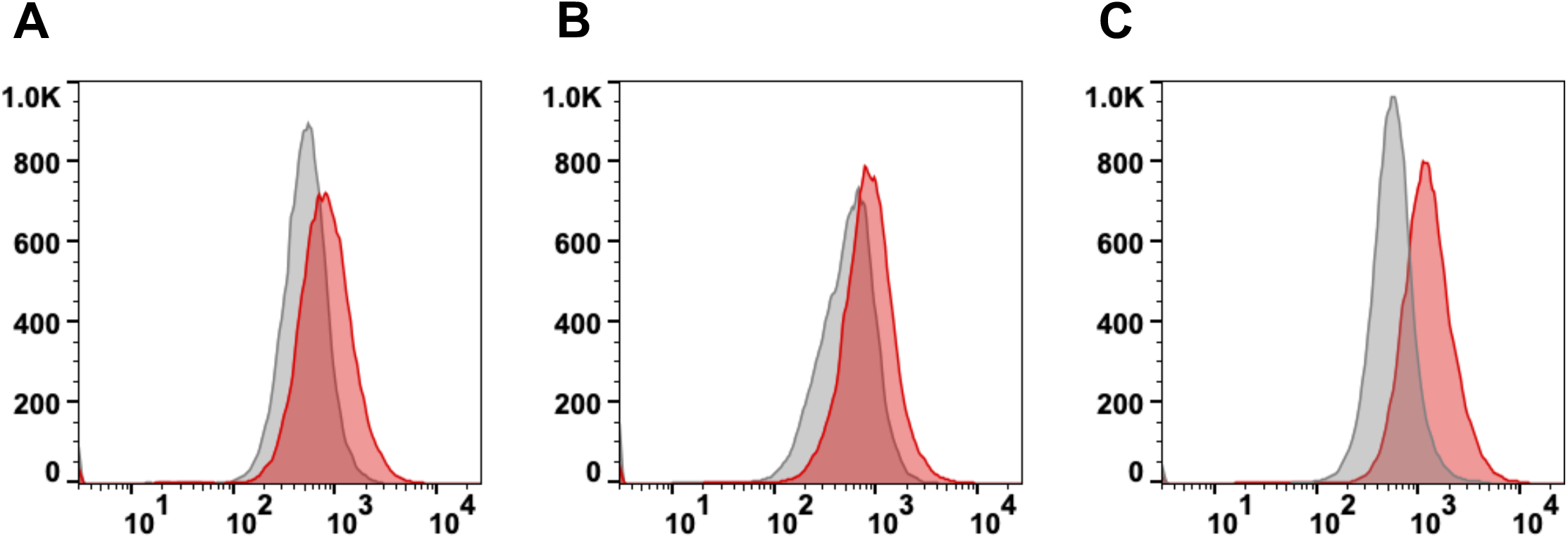
Expression of human SLAM (hSLAM) on Vero ΔPVR1/2 hSLAM+ cells. Two cell clones of Vero ΔPVR1/2 hSLAM+ cells [(A) clone 1: c1, (B) clone 2: c2] and Vero/hSLAM cells [control (C)] were stained with FITC-labeled mouse anti-human SLAM (CD150) IgG (red line) or FITC-labeled mouse control IgG (gray line).

**Figure 2.**
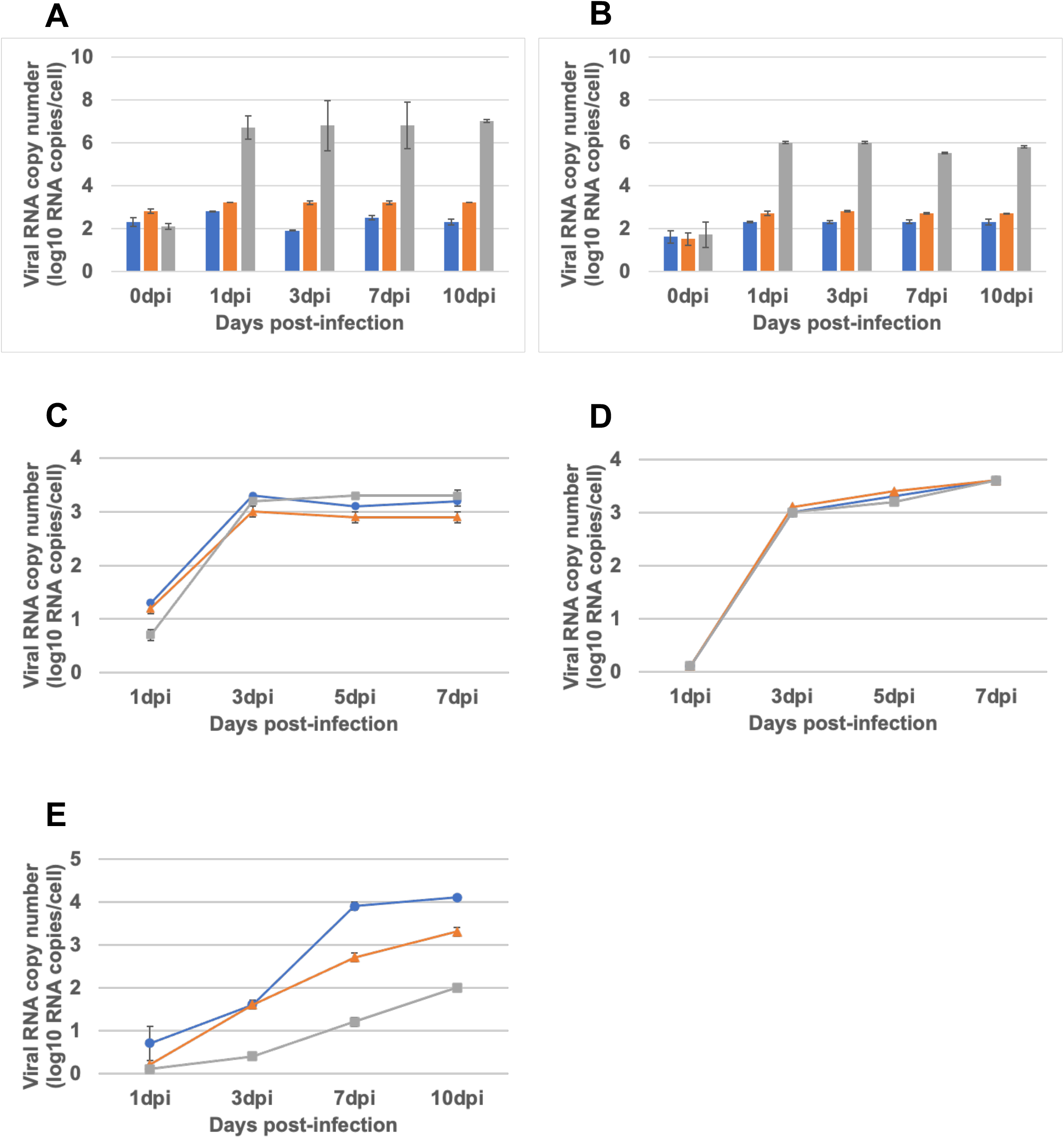
Replication of poliovirus (PV), measles virus (MeV) and rubella virus (RuV) in two cell clones of Vero ΔPVR1/2 hSLAM+ cells and Vero/hSLAM cells. (A) PV Sabin 1 infection at a MOI of 10. (B) P Sabin 3 infection at a MOI of 10. (A, B) Blue, orange, and gray bars show data for Vero ΔPVR1/2 hSLAM+ c1, Vero ΔPVR1/2 hSLAM+ c2, an Vero/hSLAM cells, respectively. (C) Wild-type MeV IC-323-EGFP infection at a MOI of 0.01. (D) Vaccine type MeV AIK-C-EGFP infection at a MOI of 0.01. (E) Wild-type RuV rHS717AG1 infection at a MOI of 0.01. Viral RNA loads were measured at the indicated time points. (C, D, E) Blue, orange, and gray graphs show data for Vero ΔPVR1/2 hSLAM+ c1, Vero ΔPVR1/2 hSLAM+ c2, an Vero/hSLAM cells, respectively.

We next conducted MeV and RuV isolation from clinical specimens. MeV specimens (10 throat swabs) were collected from measles patients in Osaka prefecture, Japan, in 2019. The infectivity titers of the MeV clinical specimens were measured by plaque assays using Vero/hSLAM cells and the two Vero ΔPVR1/2 hSLAM+ clones, as performed in a previous study (5). The viral titers measured using the two clones were comparable with those measured using Vero/SLAM cells (Table 1). These results showed that the two cell clones had the same level of susceptibility to MeV isolation as Vero/hSLAM cells. A total of 30 RuV specimens were collected from rubella patients in Osaka prefecture, Japan, in 2018 (1 throat swab) and 2019 (9 throat swabs), Aichi prefecture, Japan, in 2018 (10 throat swabs), and Toyama prefecture, Japan, in 2018 (6 throat swabs, 2 urine samples) and 2019 (2 throat swabs). These specimens were inoculated into the cells (Vero/hSLAM, Vero ΔPVR1/2 hSLAM+ c1 and Vero ΔPVR1/2 hSLAM+ c2 cells) and incubated for 7 days. Cell suspensions were harvested and the 7-day culture was repeated twice more. Replication of RuV was confirmed by a real-time PCR assy. The RuV isolation rate using Vero ΔPVR1/2 hSLAM+ c1 cells was similar to that using Vero/hSLAM cells, but was higher using Vero ΔPVR1/2 hSLAM+ c2 cells than Vero/hSLAM cells (Table 2). The RuV RNA levels in the two Vero ΔPVR1/2 hSLAM+ cell clones were greater than those in Vero/hSLAM cells. Therefore, these Vero ΔPVR1/2 hSLAM+ cell clones are advantageous for RuV isolation from clinical specimens compared with Vero/hSLAM cells.

**Table 1.** MeV titer determination for Vero ΔPVR1/2 hSLAM cells

| Throat swab<br>No. | MeV titer (PFU/ml) |  |  |
| --- | --- | --- | --- |
| | Vero ΔPVR1/2<br>hSLAM+ c1 | Vero ΔPVR1/2<br>hSLAM+ c2 | Vero/SLAM |
| 1 | $6.6 \times 10^2$ | $7.6 \times 10^2$ | $1.6 \times 10^2$ |
| 2 | $5.7 \times 10^2$ | $6.3 \times 10^2$ | $6.6 \times 10^2$ |
| 3 | - | $3.1 \times 10^1$ | $3.1 \times 10^1$ |
| 4 | $6.3 \times 10^2$ | $3.5 \times 10^2$ | $3.1 \times 10^1$ |
| 5 | $3.2 \times 10^2$ | $8.8 \times 10^2$ | $6.3 \times 10^2$ |
| 6 | $5.3 \times 10^3$ | $5.6 \times 10^3$ | $4.7 \times 10^3$ |
| 7 | $3.2 \times 10^1$ | $9.5 \times 10^1$ | $3.1 \times 10^1$ |
| 8 | -<sup>a</sup> | - | - |
| 9 | $3.8 \times 10^3$ | $4.9 \times 10^3$ | $2.9 \times 10^3$ |
| 10 | $3.8 \times 10^2$ | $9.2 \times 10^2$ | $3.8 \times 10^2$ |

a -: Not detected

**Table 2.** Rubella virus isolation from rubella patients

| Prefecture | Specimens<sup>a</sup> | RNA copy number/cell<sup>b</sup> |  |  |
| --- | --- | --- | --- | --- |
|  |  | Vero ΔPVR1/2<br>hSLAM+<br>c1 | Vero ΔPVR1/2<br>hSLAM+<br>c2 | Vero/SLAM |
| Osaka | TS-1 | - | 446±26 | - |
|  | TS-2 | - | - | - |
|  | TS-3 | - | 23±2 | - |
|  | TS-4 | 1356±83 | 1246±166 | 844±60 |
|  | TS-5 | - | - | - |
|  | TS-6 | 3226±831 | 1269±118 | - |
|  | TS-7 | - | - | - |
|  | TS-8 | 1753±345 | - | 492±25 |
|  | TS-9 | - | 1±0 | - |
|  | TS-10<sup>b</sup> | - | 507±7 | 447±26 |
| Aichi | TS-1 | - | - | - |
|  | TS-2 | - | - | - |
|  | TS-3 | - | 1797±115 | - |
|  | TS-4 | - | - | - |
|  | TS-5 | - | - | - |
|  | TS-6 | - | - | - |
|  | TS-7 | - | - | - |
|  | TS-8 | - | - | - |
|  | TS-9 | 1797±115 | 925±33 | 275±12 |
|  | TS-10 | 2761±172 | 1378±451 | 397±63 |
| Toyama | TS-1 | 2274±266 | 1890±500 | 962±291 |
|  | UR-1 | - | - | - |
|  | TS-2 | - | - | - |
|  | TS-3 | 1259±149 | 1356±372 | 412±57 |
|  | TS-4 | - | - | - |
|  | TS-5 | - | - | - |
|  | UR-2 | - | - | - |
|  | TS-6 | 1551±151 | 2405±384 | 1266±389 |
|  | TS-7 | - | - | - |
|  | TS-8 | - | - | - |

In conclusion, we have established new cell lines that are useful for the isolation of MeV and RuV, especially given the global move toward PV eradication, because the cells enable the efficient isolation and propagation of MeV and RuV from clinical specimens without concern regarding PV replication. Currently, the Japanese Collection of Research Bioresources (JCRB) Cell Bank is quality-checking these cell lines and preparing sufficient stocks for distribution in support of the global effort on the elimination of measles and rubella, and polio eradication.

## Ethics statement

The isolation of virus from clinical specimens was approved by the Medical Research Ethics Committee of the National Institute of Infectious Diseases (NIID) for the use of human subjects (approved ID: 1305).

## Acknowledgement

We greatly appreciate the staff of the JCRB cell bank at the National Institutes of Biomedical Innovation, Health and Nutrition. This work was partly supported by the Japan Agency for Medical Research and Development (AMED) under grant number JP22fk0108628.

## List of Abbreviations

PV: poliovirus
Mev: measles virus
RuV: rubella virus
PVR: poliovirus receptor
SLAM: signaling lymphocytic activation molecule
M O I: multiplicity of infection

## Notes

The authors declare no conflicts of interest.

### Competing Interest Statement

The authors have declared no competing interest.

